# Decoding Phospho-Regulation and Flanking Regions in Autophagy-Associated Short Linear Motifs: A Case Study of Optineurin-LC3B Interaction

**DOI:** 10.1101/2023.09.30.560296

**Authors:** Oana N. Antonescu, Mattia Utichi, Valentina Sora, Matteo Tiberti, Emiliano Maiani, Matteo Lambrughi, Elena Papaleo

## Abstract

Short Linear Motifs (SLiMs) play a pivotal role in mediating interactions between intrinsically disordered proteins and their binding partners. SLiMs exhibit sequence degeneracy and undergo regulation through post-translational modifications, including phosphorylation. The flanking regions surrounding the core motifs also exert a crucial role in shaping the modes of interaction. In this study, we aimed to integrate biomolecular simulations, in silico high-throughput mutational scans, and biophysical experiments to elucidate the structural details of phospho-regulation in a class of SLiMs crucial for autophagy, known as LC3 interacting regions (LIRs). As a case study, we investigated the interaction between optineurin and LC3B. Optineurin LIR perfectly exemplify a class of LIR where there is a complex interplay of different phosphorylations and a N-terminal helical flanking region to be disentangled. Our work unveils the unexplored role of the N-terminal flanking region upstream of the LIR core motif in contributing to the interaction interface. The results offer an atom-level perspective on the structural mechanisms and conformational alterations induced by phosphorylation in optineurin and LC3B recognition, along with of effects of mutations on the background of the phosphorylated form of the protein. Additionally, we assessed the impact of disease-related mutations on optineurin, accounting for different functional features.

Notably, we established an approach based on Microfluidic Diffusional Sizing as a novel method to investigate the binding affinity of SLiMs to target proteins, enabling precise measurements of the dissociation constant for a selection of variants identified in the in silico mutational screening. Overall, our work provides a versatile toolkit to characterize other LIR-containing proteins and their modulation by phosphorylation or other phospho-regulated SLiMs, thereby advancing the understanding of important cellular processes.

## Introduction

Several proteins harbor regions that are intrinsically disordered^1^, which support networks of protein-protein interactions^2,3^. Intrinsically disordered proteins or regions feature diverse interaction modes with their binding partners, including folding-upon-binding, conformational selection, formation of dynamic complexes, retained disorder in the bound state or a combination of different mechanisms at the same time ^4–8^.

Thanks to their pliability, intrinsically disordered proteins are often implicated in multiple biological processes^2,9^. Furthermore, post-translational modifications can alter the interactome of disordered proteins^10–13^.

Disordered proteins can rely on short linear motifs (SLiMs) to mediate interactions with their biological partners^4,14–16^. SLiMs are short and degenerated sequence stretches formed by few highly conserved residues (i.e., the SLIM core motif) interspersed with less conserved positions^15^. Regions flanking the core motifs and post-translational modifications, such as phosphorylation, can play a crucial role in modulating binding specificity and affinity ^17–20^. The contributions of flanking regions and phosphorylations to the binding mechanisms of SLiMs are often challenging to study both experimentally and computationally due to the transient nature of the interactions. Structure-based methods have the potential to provide an accurate understanding of SLiM modulation by phosphorylation or flanking regions, accompanied by a mechanistic understanding of their interactions with biological partners^19,21^.

SLiMs involved in the cellular clearance mechanism known as autophagy represent an intriguing case study due to their central role in the interaction with autophagy receptors and adaptors and a large variety of binding modes^19,22,23^. The ATG8 (LC3/GABARAP) family of proteins has a key role in both non-selective and selective degradation through autophagy. The LC3/GABARAP proteins interact with other proteins through the recognition of a SLiM, i.e., the LC3-interacting region (LIR), which binds to two main hydrophobic pockets (HP) of LC3 or GABARAP structures, known as HP1 and HP2 pockets^19,22,23^. The LIR motif is often associated with autophagy receptors that mediate the simultaneous binding of components of the autophagy machinery and cytosolic cargo ^24–26^, therefore representing a hub in the autophagy process. LIR amino acid sequences vary between proteins and share a core motif [W/F/Y]xx[L/I/V], whereas other residues of the LIR could contribute to the specificity toward different LC3/GABARAP family members^27–29^.

Among the known LIR-containing proteins, optineurin (OPTN) acts as a selective autophagy receptor and adaptor^30–32^. OPTN also regulates several other cellular processes associated with the trafficking of membrane-delivered cargo^30^, a trait common to scaffolding proteins. OPTN is a multi-domain protein (**Figure 1A**) and it harbors a LIR motif (**Figure 1B**), which mediates the interaction with LC3 and GABARAP proteins. The LIR motif allows OPTN to interact with LC3 and GABARAP proteins^33,34^. The LIR motif of OPTN has a role in mitophagy amplification rather than in selectivity, suggesting a general mechanism in which LC3 proteins recruit autophagy factors to drive the autophagosome growth and amplify selective autophagy in an LC3-dependent feedback loop^35^. OPTN mutations or level alterations are also associated with several different pathologies, including glaucoma, amyotrophic lateral sclerosis, Paget’s disease of bone, Crohn’s disease and cancer ^36–38^.

**Figure 1.**
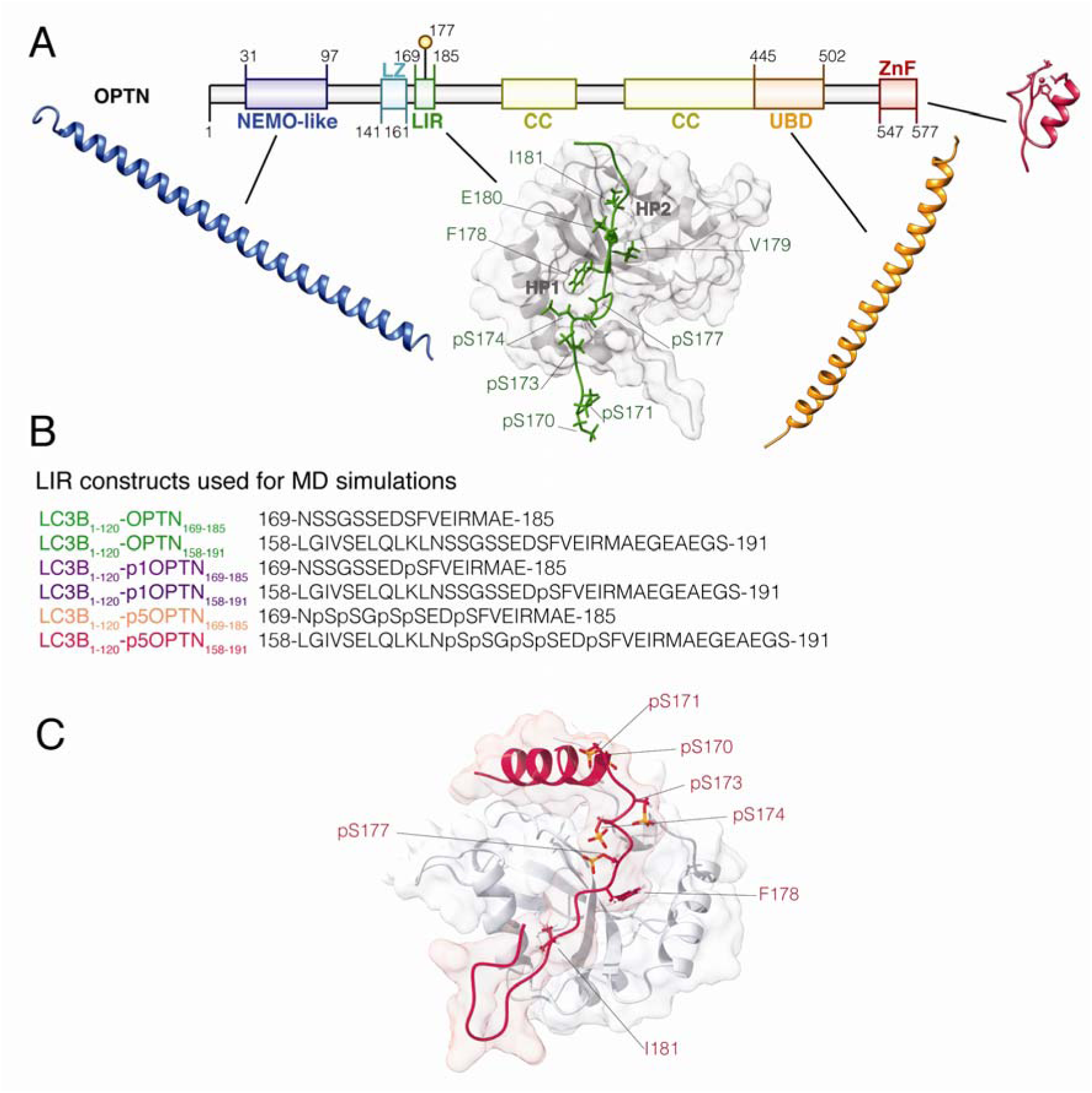
OPTN LIR region and constructs used in the molecular dynamics (MD) simulations. The figure illustrates the location of the LIR motif of OPTN and the hydrophobic residues for binding to the HP1 and HP2 pockets of LC3B. We also summarized the constructs used for the different MD simulations. The list of simulations collected in the study can be found in Table S1.

In light of the above observations, it is clear that the OPTN function needs to be tightly orchestrated and modulated at the post-translational level. In this context, OPTN undergoes phosphorylation, and a subset of these phosphorylations are relevant in the selective autophagy process known as xenophagy, namely the degradation of intracellular pathogens^36^. In particular, OPTN is phosphorylated by TANK-binding kinase 1 (TBK1) on S177 in its LIR domain, which results in its increased binding affinity to LC3B, enhancing the autophagy-mediated clearance of ubiquitin-coated *Salmonella enterica*^33,34^.

We here show how Alphafold-based models for interaction^39,40^, biomolecular simulations ^41–43^ and other structure-based computational approaches^44–47^, together with experimental biophysical data based on Microfluidic Diffusional Sizing^48,49^ are effective to study the modulation by phosphorylation of SLiMs focusing on the OPTN_LIR_-LC3B complex and its disease-related mutations. In a broader context, our work provides a solid integrative computational and experimental framework to apply to other LIRs or SLiMs regulated by phosphorylation.

## Materials and Methods

### Modeling of constructs for MD simulations

We used the NMR structure of OPTN LIR (residues 169-185, NpSpSGpSpSEDpSFVEIRMAE) in its pentaphosphorylated form (p5) in complex with LC3B (residues 1-119, PDB entry 2LUE^34^, first conformer) as a starting structure for four MD replicates of the LC3B_1-119_-p5OPTN_168-185_ complex. The structure includes an N-terminal expression tag (1-GAMG-4). Thus, we mutated it into the canonical sequence of LC3B from UniProtKB (1-MPSE-4) and added the missing C-terminal residue G120 (LC3B_1-120_-p5OPTN_169-185_). We then model its corresponding wild-type variant (LC3B_1-120_-OPTN_169-185_) by retro-mutating pS170, pS171, pS173, pS174, and pS177 to non-phosphorylated serines, using the mutagenesis tool of PyMOL (http://www.pymol.org/pymol). Furthermore, we modeled a single phosphorylated variant (LC3B_1-120_-p1OPTN_169-185_) in which we kept only pS177.

We also used AlphaFold-multimer^39^ to generate a model of the full-length complex of LC3B and OPTN and retained the first five ranked models (out of 25 models) characterized by low Predicted Aligned Error (PAE) scores. We used the AlphaFold database (version available 25/01/2023). We then retained the first ranked model to generate a trimmed version of the complex spanning the OPTN region 158-191 (LC3B1-120-OPTN158-191) and remodel the corresponding phosphorylated variants (LC3B1-120-p1OPTN158-191 and LC3B1-120-p5OPTN158-191) with ViennaPTM^50^.

### Molecular Dynamics (MD) simulations

We performed all-atom MD simulations in explicit solvent for each variant and construct (**Table S1**) using GROMACS^51^. We employed the CHARMM22* protein force field^52^, which we recently shown as a good force field for LC3B^53^. We used the TIPS3P water model^54^ and SP2 parameters for phosphorylated serine residues adapted from^55^. We used periodic boundary conditions and a dodecahedral box with a minimum distance between the protein surface and the box edges of 15 Å. We capped the N- and C-terminal groups of both LC3B (NH_3_+ and COOH, respectively) and the LIR peptides (NH_2_ and COOH). We predicted the His27, His57, and His86 tautomeric states with H++, Propka software^56^, accompanied by visual inspection. We used the Nε2-H tautomeric state for all the histidine residues in the simulation. We equilibrated the systems before the production run minimization, solvent equilibration, pressurization, and thermalization. We performed productive MD simulations at 298 K with the canonical ensemble, mimicking a concentration of 150 mM NaCl and including counterions to neutralize the system.

The MD simulations of LC3B_1-119_-p5OPTN_169-185_ were used to compare the MD results with available NMR data to evaluate the sampling quality achieved with the selected force field since they included the same construct used for NMR. The simulations with LC3B_1-120_-OPTN_158-191_ and LC3B_1-120_-OPTN_168-185_ were used for the analyses to understand the mechanisms induced by phosphorylations or the impact of disease-related mutations.

During pre-processing of the trajectories, we verified that the minimal distance between each protein complex and its image was sufficient (i.e., higher than 15 Å) to rule out artifacts due to contacts between the protein and the corresponding image in the periodic boundary conditions. We carried out the simulations without embedding LC3B in a lipidic membrane considering our previous study where we showed that the majority of the properties of interest were similar comparing MD simulations of LC3B in solution or associated with the membrane^53^. The details on the constructs used for MD simulations are reported in **Figure 1** and **Table S1**.

### Enhanced sampling simulations

We carried parallel bias metadynamics simulations for the LC3B_1-120_-p5OPTN_158-191_ variants using as starting structures the same used for the unbiased MD replicate 1 and the salt concentration and temperature. The metadynamics simulations were carried out using GROMACS and PLUMED ^51^ ^57^. We collected two different sets of simulations, including different collective variables (**Table S1**). In one case, we included two different collective variables describing the conformational changes of F178 of OPTN in the HP1 pocket of LC3B. They included the χ*_1_* sidechain dihedral of F178 and the distance between Cζ side chain atoms of OPTN F178 and LC3B F108 side chains. Moreover, we monitored during the simulation the distance between Cζ F178 and the Cδ atom of LC3B I23. In the other set of simulations, we included a parabeta collective variable to describe the intermolecular β–sheet between OPTN and LC3B. We used a bias factor of 4 (flipping mechanism) and 8 (β–sheet formation) and an initial sigma of 0.1 for all the collective variables. In all the simulations, we included an upper wall at a distance of 20 Å for the distance between the Cζ atom of F178 and the Cδ atom of LC3B I23 to avoid to sample unbinding events for the peptide. Moreover, we included a lower wall at a distance of 8 Å for the distance between the N- and C-terminal atoms of the peptide, to avoid the sampling of conformations that are too bent and would not be possible to explore in the context of the full-length protein. We carried out a block analysis for the estimate of the error associated with the calculated free energy for each simulation and each collective variable. We also calculated the differences in free energy between the minima as a function of the simulation time to evaluate convergence.

### Comparison of MD replicates with NMR data

We selected 1000 frames (equally spaced in time) for the replicate1 of the LC3B_1-119_-p5OPTN_169-185_ to predict chemical shifts from the MD simulations and compare them to the corresponding experimental data (Biological Magnetic Resonance Bank (BMRB), entry 18518). To this goal, we applied the delta_CS Python-based pipeline^58^ (https://github.com/ELELAB/delta_cs). This includes the calculation of backbone, β-carbon chemical shifts, along with chemical shifts from side-chain hydrogens for the MD ensemble with PPM ^59^ and the associated squared errors between the predicted and the corresponding experimentally measured chemical shifts. Furthermore, the pipeline includes CH3Shift^60^ to calculate chemical shifts of side-chain methyl carbon atoms (i.e., 13C shifts). It then compares them to the experimental side-chain chemical shifts. We used a reduced χ^2^ (χ^2^) metric as recently applied to another case study^61^. The square deviation between the calculated and experimental values was normalized by the variance of the chemical shift predictor for each type of chemical shift and the total number of chemical shifts. Therefore, low χ^2^_red_values indicate a good agreement between experimental and predicted chemical shifts.

### Protein flexibility

We calculated the average Root Mean Square Fluctuation (RMSF) for each Cα atom in the central part of the LIR peptides (residues 170-184). We did the calculations averaging the RMSF over 10-ns or 100-ns time windows as a probe of the LIR flexibility in the LC3B LIR-binding pocket and estimated the associated standard errors.

### Solvent-accessible surface

We calculated the side-chain solvent-accessible surface area (SASA) for the side-chain atoms of the residues of the LIR core motifs that bind to HP1 (F178) or HP2 (I181) pockets of LC3B. We estimated the SASA for the LC3B residues belonging to HP1 (D19, I23, P32, I34, K51, L53, and F108) and HP2 (F52, V54, P55, L63, I66, and I67) pockets^19^ from the MD simulations of the different variants. We also included in the analysis the LDK tripeptide (L47, D48, and K49) and the ubiquitin patch of LC3B (R10, R11, K49, and K50), which are located in the proximity of the HP1 pocket and contribute LIR-specific interactions^19,25^. We used NACCESS^62^ to calculate the relative SASA for each residue using frames extracted every 20 ps from the MD trajectories.

### Secondary structure propensity of the LIR peptides

We employed the DSSP dictionary^63^ to estimate the different classes of secondary structures for each residue of the OPTN peptides in the MD simulations. DSSP defines eight classes of secondary structure elements based on an estimate of patterns of hydrogen bonds: 3.10 helices, α-helices, π-helices, β-sheets, β-bridges, turns, bends, and coil. For each residue of interest, we calculated the occurrence of each structure class in the MD simulation.

### Analysis of electrostatic interactions and hydrogen bonds

We used PyInteraph2^64^ to calculate electrostatic interactions in the form of hydrogen bonds and salt bridges. For salt bridges, all the distances between atom pairs belonging to charged moieties of two oppositely charged residues were calculated, and the charged moieties were considered as interacting if at least one pair of atoms was found at a distance shorter than 5 Å. We used the atoms forming the carboxylic group for aspartate and glutamate. The NH3 and the guanidinium groups were employed for lysine and arginine, respectively. A hydrogen bond was identified when the distance between the acceptor atom and the hydrogen atom was lower than 3.5 Å, and the donor-hydrogen-acceptor atom angle was greater than 120°. We calculated main-chain main-chain, main-chain side-chain, and side-chain side-chain hydrogen bonds. We customized the configuration files used by PyInteraph2 for the identification of salt bridges and hydrogen bonds by adding the oxygens of the phosphate group to the charged groups (for salt bridges) and to the hydrogen bond acceptors (for hydrogen bonds) to include the phosphorylated serines. We retained only intermolecular interactions present in at least 20% of the simulation frames (*pcrit* = 20%), as previously applied to other proteins^65–68^.

### Atomic contact-based Protein Structure Network (PSN)

We used the PyInteraph2 implementation^64^ of the PSN method based on atomic contacts as originally formulated by Vishveshwara’s group^47^ and applied to the MD ensemble of the different constructs used in this study. We retained the pairs of residues whose sequence distance is higher than 1 for the edge calculations and at a distance lower than 4.5 Å. In the calculation, normalization factors for each residue are applied as previously discussed^69^ to calculate the interaction strength. We used an I_min_ value of 3. For phospho-serines, we used the normalization factor reported in a previous paper^70^. In addition, we retained edges with occurrences higher than 50% of the frames in the MD ensemble in the final PSN. The interaction strength associated with each edge in the final PSN corresponds to the average interaction strength for the edge over all the ensemble structures, and it is used to weight the PSN.

### Changes in binding free energy

We employed Foldx5 ^44^ to perform *in silico* saturation mutagenesis and estimate changes in binding free energy upon mutations using the Python wrapper MutateX^71^. We estimated the average difference in binding free energy (ΔΔGs) upon mutations using 25 representative structures from the MD simulations of LC3B_1-120_-OPTN_158-191_, LC3B_1-120_-p1OPTN_158-191,_ and LC3B_1-120_-p5OPTN_158-191_. This strategy may help mitigate the limitation due to the backbone stiffness of FoldX^71,72^. As a reference, we also performed similar calculations using as starting structure the 20 NMR-derived structures of LC3B_1-119_-p5OPTN_168-185_ deposited in the PDB entry 2LUE). We calculated the average ΔΔGs for each position, and we used a cutoff of 1kcal/mol in ΔΔG to classify the effects of the mutations^71^. In particular, we defined stabilizing or destabilizing variants featuring ΔΔG values lower than -1 or higher than 1 kcal/mol, respectively.

### MAVISp analyses for disease-related variants of OPTN_158-191_

We used the data deposited in the MAVISp database^73^ where variants found in COSMIC^74^, cBioPortal^75^, and ClinVar^76^ have been retrieved for OPTN (Uniprot ID Q96CV9, and RefSeq ID NP_068815).

We retained only mutations in the region used for the modeling, i.e., 158-191 (i.e., V161M, E163D, L166V, N169S/T, E175Q, D176Y, F178V, E187Q, V179D, A184D/S). We then applied the MAVISp framework ^73^ for the assessment of the effects of these variants on: i) structural stability, ii) interactions with LC3B, iii) homodimerization, iv) interplay with post-translational modifications and v) long-range effects.

We used the model downloaded from the AlphaFold database^77^ on the 23rd of March 2023 for the STABILITY module of MAVISp. We used the starting structure for the simulations of the non-phosphorylated variant (LC3B_1-120_-OPTN_158-191_), and 25 representative frames from the MD replicate 1 for the LOCAL INTERACTION module in the simple and ensemble mode, respectively. We also used a model generated with AlphaFold multimer and experimentally validated^78^ to assess the effects of the mutations on the homodimerization of OPTN_1-577_.

In addition, we retrieved (REVEL, AlphaMissense) or calculated (DeMask, EVE) the pathogenicity scores associated with the mutations using REVEL^79^, DeMask^80^, EVE^81^ and AlphaMissense^82^ with MAVISp workflows.

### Structural clustering

We calculated pairwise RMSD values for each trajectory using the main-chain atoms of the complex. We performed clustering on the RMSD matrix using the GROMOS algorithm^83^ and an RMSD cutoff equal to the average RMSD from each MD replicate. We then retrieved the average structures for the first three populated clusters for the simulations of the unphosphorylated variant LC3B_1-120_-OPTN_158-191_. The average structure of each cluster corresponds to the structure belonging to the cluster, which features the lowest RMSD values to its neighbors in the cluster. These have been used as starting structures for the calculations in the ensemble mode with MAVISp and the LOCAL_INTERACTION module with the flexddg protocol of RosettaDDGPrediction (see above).

### Correlogram

We collected correlation statistics on the MD replicates to investigate the relationship between selected pairwise distances between representative side-chain atoms of the residues involved in salt bridges. To perform this analysis, we utilized R script adapted from our previous work ^84^. We measured linear dependencies using the Pearson correlation coefficient as a metric and we calculated the associated p-values. For visual representation, we constructed the correlograms with a hierarchical clustering approach to rank the measurements based on the strength of their correlation and used a significance level corresponding to a p-value of 0.01.

### Protein purification

The construct encoding for GST-fused LC3B was expressed in *E. coli* BL21 (DE3) cells, and the expression of the fusion proteins was induced by adding 0.5 mM IPTG at 37[[°C for 4 hours. Cells were lysed by sonication in lysis buffer (50 mM Tris-HCl pH 8, 250 mM NaCl, 1 mM DTT, cOmplete™, EDTA-free Protease Inhibitor Cocktail tablets, and PhoSTOP phosphatase inhibitor tablets). Pierce™ Glutathione Magnetic Agarose beads were used to precipitate the GST-fused protein for 2 hours at room temperature under shaking. After washing of the beads in lysis buffer, the protein was eluted in elution buffer (50 mM Tris-HCl pH 8, 10 mM reduced glutathione). The buffer was exchanged in assay buffer (10 mM Tris-HCl pH 7.4, 150 mM NaCl, 0.1 % Tween-20) using PD10 gravity columns (GE Healthcare). The protein was concentrated using Amicon Ultra-0.5 10K Centrifugal Filter Devices (Merck Millipore). The protein concentration was measured on a Nanodrop 1000 spectrophotometer at 280 nm using the extinction coefficient of Trp and Tyr residues in the GST-LC3B sequence (48820 cm-1M-1).

### Peptides

N-terminally labeled FITC-Ahx OPTN LIR peptides were purchased from TAGCopenhagen in >95% purity. We have used the following peptide sequences corresponding to the 169-185 residues in full-length OPTN:

FITC-Ahx-NSSGSSEDSFVEIRMAE (WT),

FITC-Ahx-NSSGSSEDpSFVEIRMAE (pS177),

FITC-Ahx-NSSGSSEDDFVEIRMAE (S177D), FITC-Ahx-NSSGSSEDEFVEIRMAE (S177E),

FITC-Ahx-NSSGSSEDSWVEIRMAE (F178W),

FITC-Ahx-NSSGSSEDpSWVEIRMAE (pS177 F178W),

FITC-Ahx-NSSGSSEDpSFVEIRAAE (M183A), and FITC-Ahx-NSSGSSEDpSAVEIRMAE (F178A),

where “pS” is phosphorylated Ser.

Sequences were confirmed by mass spectrometry and purity by RP-HPLC. The peptides were dissolved in an assay buffer and their concentration was measured on a Nanodrop 1000 spectrophotometer at 280 nm using the extinction coefficient of the FITC group (18542 cm-1M-1) and Trp residue where present.

### Microfluidic Diffusional Sizing (MDS)

All MDS experiments were performed on a Fluidity One-M instrument at room temperature using commercial chips (Fluidic Analytics, Cambridge, UK). FITC-Ahx-labeled OPTN_169-185_ LIR peptides at a total concentration of 1 µM, together with a varying concentration of GST-LC3B, were added and diluted in assay buffer. The samples were incubated at room temperature for at least 2 hours to reach equilibrium. The change in hydrodynamic radius with GST-LC3B concentration was fitted with an in-house-developed Python script using a one-site total binding model, to obtain Kd values. The equation used for fitting the data was:

Y=Rh_free+(Rh_complex-Rh_free)*((X+n*L+Kd)-((X+n*L+Kd)^2-4*n*L*X)^0.5)/(2*n*L)

where:

Rh_free = Hydrodynamic radius of the labeled peptide (nm)

Rh_complex = Hydrodynamic radius of the complex between the labeled peptide and unlabeled GST-LC3B (nm)

n = binding sites occupied on the labeled species at saturation (i.e., stoichiometry) L = the labeled peptide concentration (nM)

K_d_ = the equilibrium dissociation constant or affinity of the complex (nM)

Each MDS experiment was performed in 3-5 biological replicates, each with 1-3 technical replicates. The differences in Kd, Rh_free or Rh_complex between the tested peptide variants and the pS177 or WT variant were assessed by an unpaired, parametric t-test. Where the tested variant had a variance of more than 4-fold of the pS177 or WT variant, we used the t-test with Welch’s correction.

### Surface plasmon resonance (SPR)

SPR experiments were carried out on a Biacore X100 instrument (Cytiva). The instrument was primed into running buffer (HBS-EP+ 1x buffer, Cytiva). Following standard procedures for amine coupling, 1 μM of GST-LC3B in 10 mM sodium acetate, pH 5, was used for coupling to the carboxymethylated dextran matrix of a CM5 sensor chip surface (FC2). A reference surface was treated identically, but not subjected to GST-LC3B for immobilization (FC1). These SPR chips and HBS-EP+ running buffer were used for all experiments with GST-LC3B described in this paper, and all data collected was referenced to FC1. OPTN_169-185_ peptide variants were dissolved in running buffer at concentrations between 600 nM and 300 μM. Various concentrations of all OPTN peptide constructs were injected at 30 μl/min and 25°C, with an association time of 60 s and a dissociation time of 60 s. Resonance units (RU) were recorded as a function of time. The BIAevaluation software package was used for data analysis using a simple 1:1 model.

## Results and Discussion

### Overview of the complex between OPTN and LC3B

The LIR (residues 169-185) of OPTN is located C-terminal to a coiled-coil region and in a disordered tract, according to the AlphaFold2 model deposited in the AlphaFold database (**Figure 1C**). The core region of the motif is encompassed by the sequence 178-FVEI-181, where F178 and I181 occupy the HP1 and HP2 hydrophobic pockets of LC3B, respectively (**Figure 1B**). S177 can be phosphorylated in position -1 to the phenylalanine (**Figure 1B**). Four additional serine residues (S170, S171, S173, and S174) N-terminal to the LIR core motif can also undergo phosphorylation (**Figure 1B**). The structure of the complex between an OPTN LIR variant with the five phospho-serines in complex with LC3B (LC3B_1-119_-p5OPTN_169-185_) has been solved by NMR^34^ and used as a starting structure for the modeling and simulations in this work (**Table S1**).

Moreover, we collected one-microsecond unbiased MD simulations of the unphosphorylated (LC3B_1-120_-OPTN_169-185_ and LC3B_1-120_-OPTN_158-191_), mono-phosphorylated (LC3B_1-120_-p1OPTN_169-185_ and LC3B_1-120_-p1OPTN_158-191_) and the multiple phosphorylated variants (LC3B_1-120_-p5OPTN_169-185_ and LC3B_1-120_-p5OPTN_158-191_) all without the tag sequence used for NMR or including a N-terminal flanking region which has helical propensity (**Table S1**).

### MD simulations of LC3B in complex with phosphorylated OPTN are in good agreement with NMR chemical shifts

Before proceeding with the analyses of the MD ensembles, it is essential to assess the quality of the structures collected in the simulations, i.e., the conformational sampling. Hence, we compared the MD ensembles with chemical shift data from NMR, similar to what we did for other case studies^53,58,61,85^.

For this purpose, we carried out four one-μs MD replicates of LC3B_1-119_-p5OPTN_169-185_. The NMR data of LC3B_1-119_-p5OPTN_169-185_ includes the assignment of backbone and side-chain chemical shifts^34^. Chemical shifts are probes of dynamics along different time scales^86–88^ and can also be predicted by MD simulations ensemble of structures^59,89,90^. The calculated chemical shifts from our simulations agree with the experimental data with low χ2_red_ values, which converged to a plateau after 200-300 ns (**Figure 2A**).

**Figure 2.**
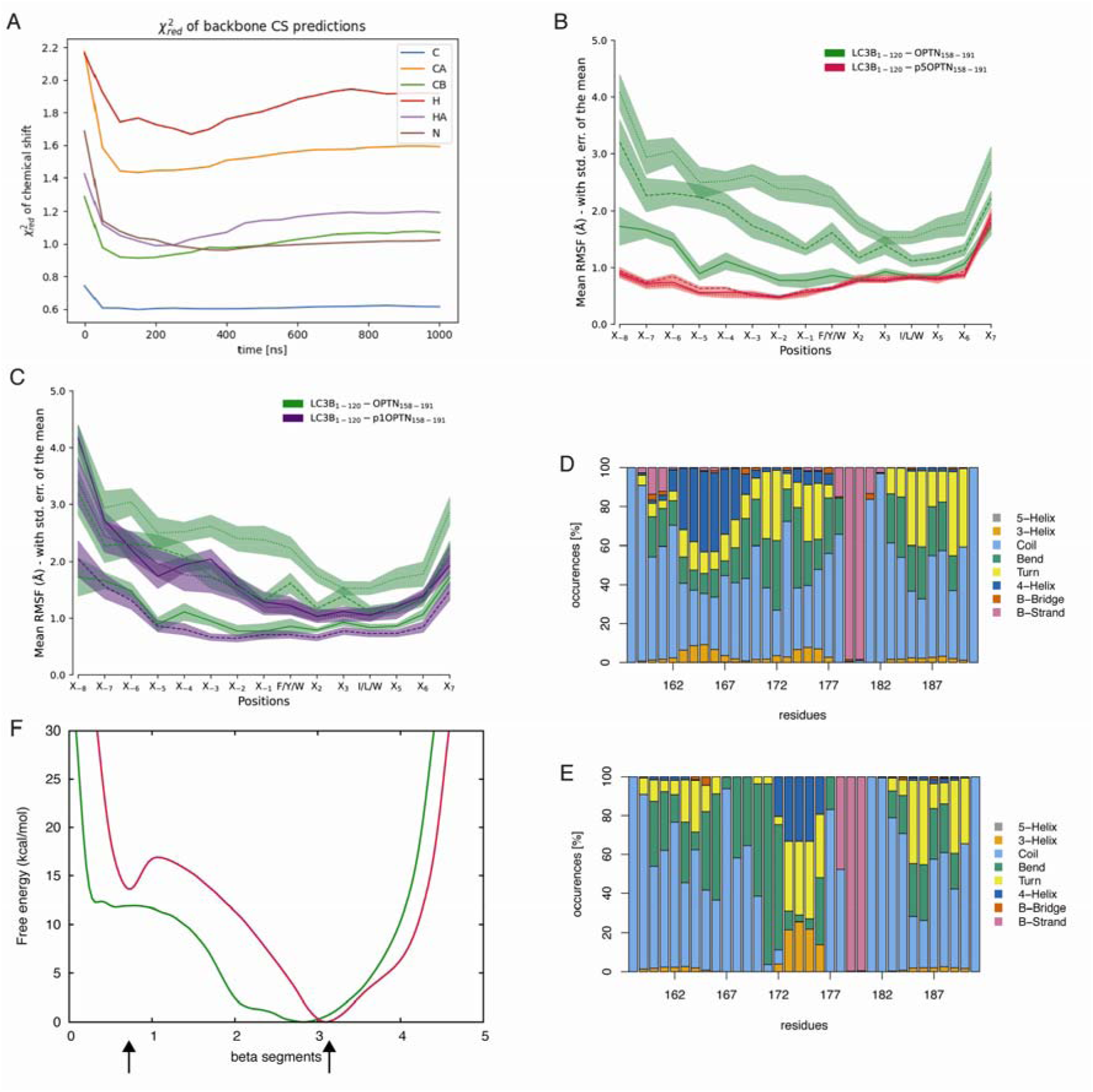
Comparison with experimental data and changes in protein flexibility and secondary structures induced by phosphorylation. **A)** For each replicate of the LC3B_1-119_-p5OPTN_169-185_ construct, we estimated a reduced χ^2^ (χ^2^_red_) over the simulation time. This value provides a measurement of the agreement between the experimental chemical shifts from NMR and the calculated chemical shifts from the MD trajectories. The metric is calculated for each atom type. Low values of χ^2^_red_ indicate better agreement between experiments and simulations. We reported the example of the MD replicate 1 of LC3B_1-119_-p5OPTN_169-185_ for the sake of clarity. **B-C)** The average Cα RMSF for each replicate, calculated using 100-ns time windows is showed for LC3B_1-120_-p5OPTN_158-191_ (B) and LC3B_1-120_-p1OPTN_158-191_ (C) compared to the unphosphorylated construct, respectively. **D-E**) The bar plots show the secondary structure content according to the DSSP dictionary using a concatenated trajectory with all the MD replicates for LC3B_1-120_-OPTN_158-191_ (D) and LC3B_1-120_-p5OPTN_158-191_ (E). It can be observed that the multiple phosphorylation slightly locally increases the propensity to β-strand formation. **F)** The panel shows the mono-dimensional free energy surface (FES) profiles for the parabeta collective variable employed in the metadynamic simulations to describe the intermolecular β–sheet between OPTN and LC3B, i.e., measuring the number of segments that resemble a parallel β-sheet conformation. The wild-type LC3B_1-120_-OPTN_158-191_ and penta-phosphorylated LC3B_1-120_-p5OPTN_158-191_ are highlighted in green and red respectively. We observe that the phosphorylation induces an increase of intermolecular β-sheet formation.

Furthermore, we calculated the side-chain chemical shifts for methyl-containing residues (Ile, Val, and Leu), which are suitable probes of local conformations (**Table 1**) and are also associated with preferences for different rotameric states of each residue in solution^91–93^ . Most of the chemical shift classes featured predicted values in agreement with the experimental data, with low χ^2^ indicating that the MD sampling capture the overall dynamics observed experimentally in solution. The only exception is isoleucine residues with a χ^2^_red_ higher than 3 for the Cδ depending on the replicate under investigation (**Table 1**).

**Table 1.** Comparison between experimental and predicted side chain chemical shifts for methyl groups. The results for the four MD replicates of LC3B_1-119_-p5OPTN_169-185_ are reported in the table. We report the reduced χ^2^ for each replicate and residue. Low χ^2^_red_ values indicate a good agreement with the experimental data.

| LC3B residue type | Replicate 1 | Replicate 2 | Replicate 3 | Replicate 4 |
| --- | --- | --- | --- | --- |
| ILE (Cδ) | 3.75 | 4.16 | 2.81 | 3.31 |
| ILE (Cγ2) | 0.73 | 0.72 | 0.56 | 0.64 |
| LEU (Cδ1) | 1.35 | 1.34 | 1.36 | 1.34 |
| LEU (Cδ2) | 0.34 | 0.36 | 0.46 | 0.42 |
| VAL(Cγ1) | 0.59 | 0.61 | 0.59 | 0.60 |
| VAL(Cγ2) | 0.29 | 0.24 | 0.39 | 0.30 |

With MD simulations of reasonable quality and in agreement with the protein dynamics in solution on the microsecond timescale, it becomes meaningful to use the MD ensembles to study the effects induced by phosphorylation and unveil new details of the underlying mechanism.

### The phosphorylation reduces the flexibility of the OPTN LIR in the binding pocket of LC3B

Phosphorylation of LIR motifs can decrease the flexibility of the LIR peptide in complex with LC3B^94^, stabilizing the conformation of the LIR residues in the binding pocket of LC3B. We thus investigated if the phosphorylation of the LIR of OPTN could exert a similar effect. We used the RMSF of the Cα atoms of the LIR peptide as a probe for changes in structural flexibility using the LC3B_1-120_-OPTN_158-191_ constructs. We observed a marked decrease in LIR flexibility upon phosphorylation of the OPTN LIR (**Figure 2B**), whereas the unphosphorylated peptide displays higher RMSF values along the entire sequence. The effect is observable at different timescales, as attested by the analyses carried out within ten and 100-ns time windows (**Figure 2B**). The phosphorylation of only S177 is not sufficient to decrease the flexibility of the LIR in the complex (**Figure 2C**).

### Phosphorylation of OPTN increases the propensity to β-strand formation in the LIR binding pocket

LIRs, like other SLiMs, can have different secondary structure propensity upon binding with an interactor. Indeed, it has been experimentally shown that several LIRs fold upon binding, inducing the formation of an intermolecular parallel β-sheet with the β2 strand of LC3B. In contrast, other LIRs retain disorder in the bound state^25,34^. We thus evaluated if the phosphorylation could affect the secondary structure propensity of the OPTN LIR in a complex with LC3B using the simulations with the LC3B_1-120_-OPTN_158-191_ constructs (**Figure 2D-E**, **Figure S1**).

We only observed the two residues downstream to F178 (i.e., 179-180) in the unphosphorylated variant with a propensity for β-strands (**Figure 2D**). The phosphorylation of S177 seems sufficient to slightly locally increase the β-strand propensity, including F178 (**Figure S1**). The multiple phosphorylation did not increase further β-strand formations with respect to the mono-phosphorylated variant (**Figure 2E, Figure S1**). The β-strand of OPTN contributes to the intermolecular β-sheet via interaction with the β-strand 2 of LC3B, as observed in the experimental structures of the multiple phosphorylated variants or of the phospho-mimetics^34^.

We thus evaluated if this behavior could be due to limited conformational sampling in the unbiased MD simulations using an enhanced sampling method based on metadynamics and parallel bias (see Materials and Methods). The collective variable used for these simulations measure the number of segments that resemble a parallel β-sheet conformation. The metadynamics results confirmed that the phosphorylation induced an increase of intermolecular β-sheet formation (**Figure 2F**).

### Phosphorylation of OPTN does not change the solvent accessibility of the core LIR residues but influences charged residues in the LDK tripeptide and in the LC3B HP1 pocket

Next, we verified if the decreased flexibility of the peptide in the pocket caused by the phosphorylation was also related to changes in solvent accessibility of the side chains of the two hydrophobic residues of the LIR core motif (i.e., F178 and I181). We could not observe marked differences in the distribution of the SASA values of HP1 and HP2 residues (**Figures S2 and S3**). Furthermore, we evaluated if the residues lining the HP1 and HP2 pockets of LC3B, the LDK tripeptide, or the LC3B ubiquitin patch were sensitive to changes in solvent accessibility. We observed a marked effect on K51 in the HP1 pocket, where the phosphorylation of S177 is sufficient to exert the effect (**Figure 3A**). Moreover, the multiple phosphorylations also decreased the solvent accessibility of D48 and K49 in the LDK tripeptide (**Figure 3B-C**). Other residues of LC3B were not influenced by changes in the accessibility of their side chains upon phosphorylation.

**Figure 3.**
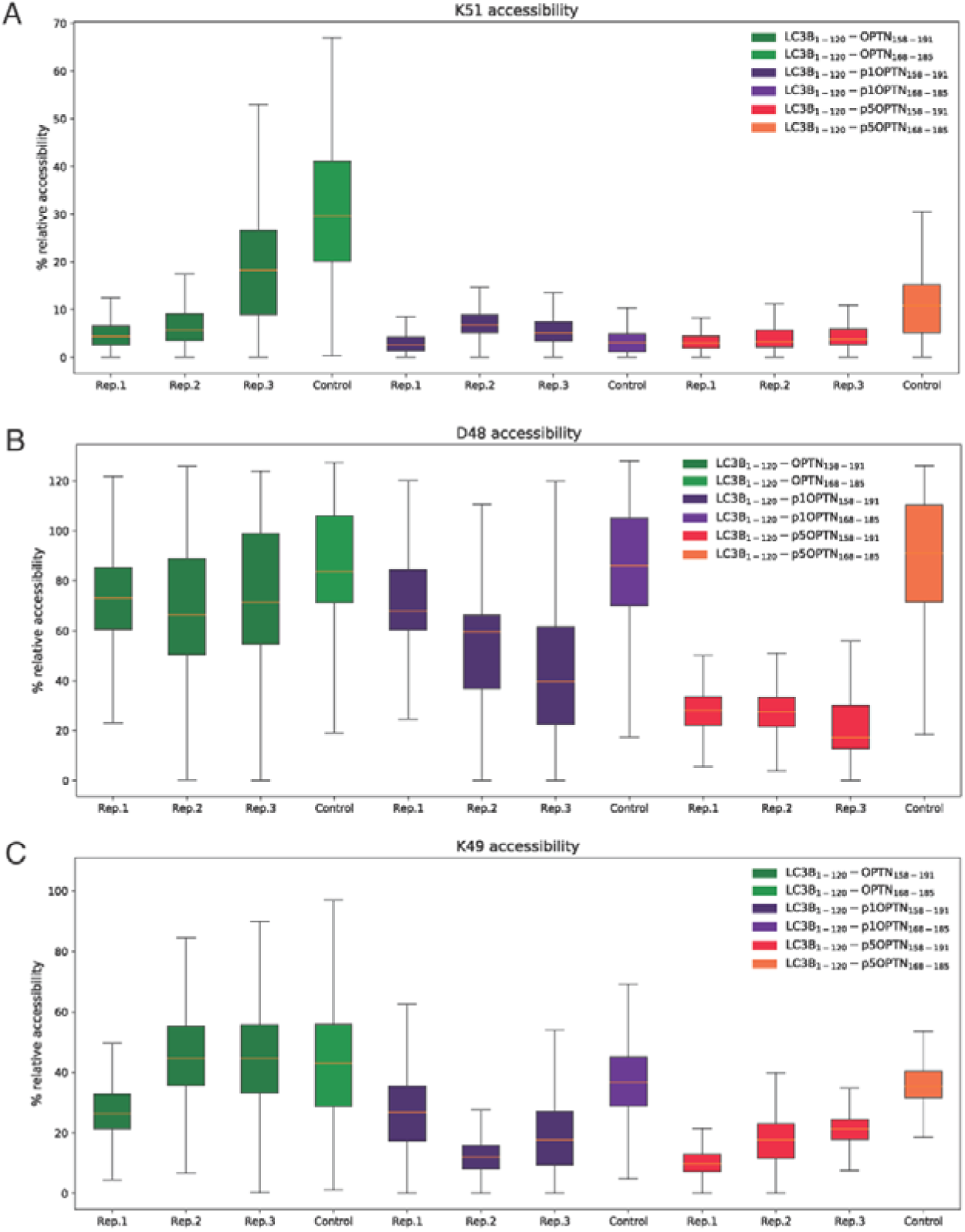
Phosphorylated OPTN_158-191_ causes changes in solvent accessibility of charged residues of LC3B at the interaction interface. The relative solvent accessibility of side chain atoms is reported for K51 (A), K49 (B), and D48 (C) in the different replicates of LC3B_1-120_-OPTN_158-191_ and LC3B_1-120_-OPTN_168-185_ variants.

In summary, the decreased structural flexibility seems not to cause changes in the residues of the LIR core motif, but it influences the accessibility of the interacting residues in LC3B.

### The phosphorylation increases the interaction strength between the LIR and LC3B both locally and long-range

Furthermore, we evaluated if the changes in solvent accessibility and flexibility were associated with changes in the interaction strength between OPTN and LC3B. We used the pairwise interaction strength as calculated from the atomic-contact-based protein structure network (see Materials and Methods, **Table S2 and** **Figure 4A**).

**Figure 4.**
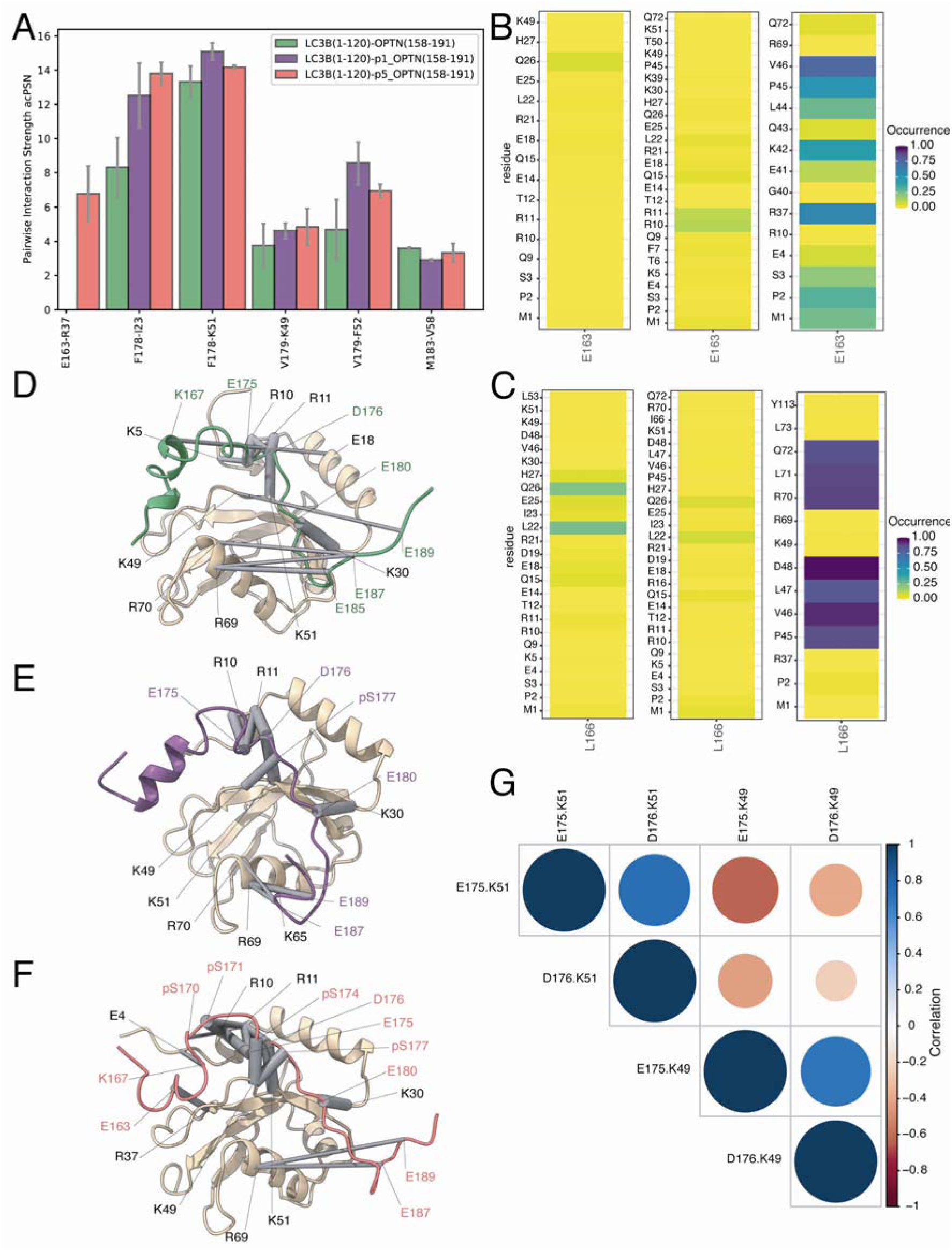
Changes in intermolecular interactions upon phosphorylation. A) The histograms report the results of pairwise Interaction Strength as calculated using an atomic-contact-based protein structure network for the pairs more sensitive to changes upon phosphorylation for the LC3B_1-120_-OPTN_158-191_, LC3B_1-120_-p5OPTN_158-191_. and LC3B_1-120_-p1OPTN_158-191_ constructs. The full set of data on interaction strengths is reported in Table S2. B-D) The networks of salt bridges are reported on the 3D structure of the first frame of the MD simulations of LC3B_1-120_-OPTN_158-191_ (B), LC3B_1-120_-p1OPTN_158-191_ (C) and LC3B_1-120_-p5OPTN_158-191_ (D) using the MD replicate 1 for sake of clarity. The salt bridges are depicted as gray sticks, whose size is proportional to their occurrence in the simulation. The full set of data is reported in Table S3.

We observed that the phosphorylation of pS177 and even more of the five serine residues increased the interaction strength between F178 of OPTN and I23 of the HP1 pocket of LC3B. Moreover, it also increased the strength of interaction between V179 of OPTN and F52 of the HP2 pocket of LC3B. In contrast, the LIR core residue I181 or the residue M183 in the C-terminal flanking region (**Figure 4A**) featured similar interaction strengths in all the simulations. The phosphorylation at multiple sites also has an additional long-range effect, increasing the interaction strength in the N-terminal flanking region of the OPTN LIR between E163 and R37 (**Figure 4A**).

This effect cannot be recapitulated by the single phosphorylation at position S177. The role of E163 as important interaction hotspot in the penta-phosphorylated variant of OPTN is also confirmed by the analyses carried out with another atomic-contact based method that allows to evaluate not only the occurrence but also the number of contact formation/breaking during the simulation time (**Figure 4B**). This analysis also disclosed K42 as additional stable intermolecular interaction for E163 upon phosphorylation. Furthermore, this second set of analysis allowed to identify L166 as additional site in the N-terminal flanking region of OPTN switched on by phosphorylation (**Figure 4C**) and interacting with other hydrophobic residues as L71, L47 and V46.

### A rewiring of intermolecular salt bridges upon phosphorylation of OPTN with both local and long-range effects

To evaluate if phosphorylation of the OPTN LIR could affect electrostatic interactions, we estimated salt bridges (**Table S3,** **Figure 4D-F**) from the MD simulations of the LC3B_1-120_-OPTN_158-191_ variants (see Materials and Methods).

The interaction between E180 of OPTN and K30 of LC3B is the only salt bridge that is conserved at high occurrence (> 90%) in all the complexes. The unphosphorylated variant has a different pattern of salt bridges with occurrence higher than 20% depending on the MD replicate under consideration, changing from only E180-K30 in replicate 3 to some more sparse and lower occurrence salt bridges involving different residues at the interaction interface in replicates 1 and 2 (**Table S3**). Interestingly, E175 and D176 of OPTN can interchange in their role of partners of interactions for K49 and K51 of OPTN in a fluid salt-bridge network, as it can also be observed by a correlation analysis of the time series of their distances (**Figure 4G**). In particular, the formation of salt bridges by E175 and D176 with K51 is negatively correlated with the interaction of the same OPTN residues with K49, suggesting that they are mutual exclusive (Figure 4G).

The single phosphorylation of S177 rewired the electrostatic interactions toward a less dynamic and heterogeneous behavior. The high-occurrence salt bridges were strengthened and concentrated in the central part of the interface with only three main players, i.e., D176, pS177, and E180, stably interacting with K51, K49, and K30, respectively (**Table S3**, **Figure 4D-F**). The interaction between K51 and pS177 agrees with what was observed with NMR experiments and changes in affinity upon a K51A mutation on shorter constructs of OPTN^34^. We observed the same rewiring of interactions for the MD simulations of the pentaphosphorylated variant. Moreover, pS170 and pS171 contributed to stabilizing and expanding the network of electrostatic interactions involving R10 and R11 or LC3B. In addition, in agreement with the long-range effects observed above, the multiple phosphorylations promoted more stable electrostatic interactions in the N-terminal region of OPTN, where E163 and K167 also participated in salt bridges, and pS174 contributed to strengthening the interaction with K49 (**Table S3,** **Figure 4B-D**). The other phospho-residues were not involved in stable salt bridges with LC3B. The results above agree and provide a mechanistic explanation for the decreased solvent accessibility of K49 and K51 in the phosphorylated variants observed in **Figure 3**.

Interestingly, K49 also engages in a salt bridge with the residue corresponding to S177 in the LIR motif of p62 (i.e., D337), suggesting that this lysine may participate in the LIR-binding process by recruiting negatively charged residues from different positions of the N-terminal region of a LIR peptide^94^.

The analysis of intermolecular hydrogen bonds involving either main chain or side chain atoms in the different MD replicates confirmed a similar rewiring of interactions (**Table S4**). In a previous study with shorter constructs for the OPTN LIR, the phosphorylated structure was studied using NMR for the phosphorylated variants. Moreover, an additional structure of a penta-phosphomimetic replacing the serines with glutamates was solved by X-ray^34^. In the X-ray structure, the authors observed an inward flip of R11 to make hydrogen bonds with the side chain of E177 of LC3B. The authors could not detect the amide proton of R11 side chain with NMR. We thus evaluated if a similar mechanism was occurring in our simulations with longer constructs for the interaction between R11 and pS177. We collected the distances over time between the side chains of the two residues in the penta-phosphorylated simulations and evaluated their probability distribution (see OSF repository). We found the two residues at distances larger than 10 Å in our simulations, whereas we noticed that R11 mostly sense another phospho-site, i.e., pS171. The mechanism suggested here could still explain the changes in the chemical shift for R11 upon phosphorylation observed with NMR. The control simulation with a shorter construct (LC3B_1-119_-p5OPTN_169-185_) featured the interaction between pS177 and R11, as observed in the model derived by the NMR data and deposited in PDB.

### Phosphorylation does not reduce the conformational changes of the F-type LIR in the aromatic residue for binding to the HP1 pocket of LC3B

OPTN has an F-type LIR, i.e., at the core position for binding to the HP1 pocket of LC3 protein the tryptophan is replaced by a phenalanine. Phenylalanine could be more prone to undergo conformational changes in the binding pocket that cause a flipping of its sidechain outside the binding pocket. We speculate that phosphorylation could be a mechanism to reduce the flipping and provide a stronger binding at the HP1 pocket. We thus used enhanced sampling simulations to address this hypothesis based on parallel bias metadynamics. The collective variables used to describe the flipping events are showed in **Figure S4**.

At first, we analyzed the unphosphorylated variant to evaluate if the flipping could occur and in which population of the conformational ensemble. We observed one favored state when the side chain of F178 is well buried in the LC3B HP1 pocket, in contact at distances lower than 5 Å with F108, and its χ*1* dihedral is either in a minus or trans state (**Figure S4**). A minor state where is in a flipped conformation of F178 outside the binding pocket where the χ*1* dihedral angle is in a plus conformation and the distance from F108 higher than 9 Å. Neither the introduction of the phosphorylation at position S177 or of the five phosphorylation could decrease the flipping of F178 outside the HP1 hydrophobic pocket of LC3B (**Figure S4**).

### High-throughput mutational scans identify binding hotspots and residues conferring specificity for the binding mode of the phosphorylated variant

The integration of MD-derived ensembles of conformations with empirical free energy calculations^53,71,72,85,95,96^ could shed light on hotspots for the binding between LIR and LC3B proteins. According to a structure-based framework that we recently developed, i.e., MAVISp framework ^73^, we rely on a consensus approach based on FoldX and Rosetta. However, the protocols for Rosetta-based free energy calculations do not support phospho-residues, and one should rely on phosphomimetics. To determine the effectiveness of phosphomimetics in replicating the effects of phosphorylation, we used synthesized OPTN LIR peptides, introducing a phosphoserine at position 177 (pS177), as well as phosphomimicking substitutions (S177D and S177E). Subsequently, we conducted MDS experiments to evaluate the binding affinity of these peptides to GST-LC3B (**Figure 5A**). The affinity of the pS177 peptide was measured at 14.26 µM (**Figure 5A1**), while the S177D and S177E variants exhibited affinities of 44.81 µM (**Figure 5A2**) and 42.82 µM (**Figure 5A3**), respectively. The difference between pS177 and S177D does not seem significant (p = 0.123), possibly due to the high variability of the S177D data, while the difference between pS177 and S177E is statistically significant (p = 0.001). Existing literature data indicates that the affinity of the wild-type peptide falls within the range of 40-60 µM^33,34^. Therefore, it appears that the single phosphomimetic mutation at position 177 might confer a modest increase in binding affinity compared to the wild-type sequence. Still, the mimetics do not reach the level observed with authentic phosphorylation. Consequently, we decided not to include Rosetta-based calculations in this study and relied on FoldX calculations, using 25 representative structures from each of the MD replicates of the constructs LC3B_1-120_-OPTN_169-185_ and LC3B_1-120_-OPTN_158-191_ from **Table S1**.

**Figure 5.**
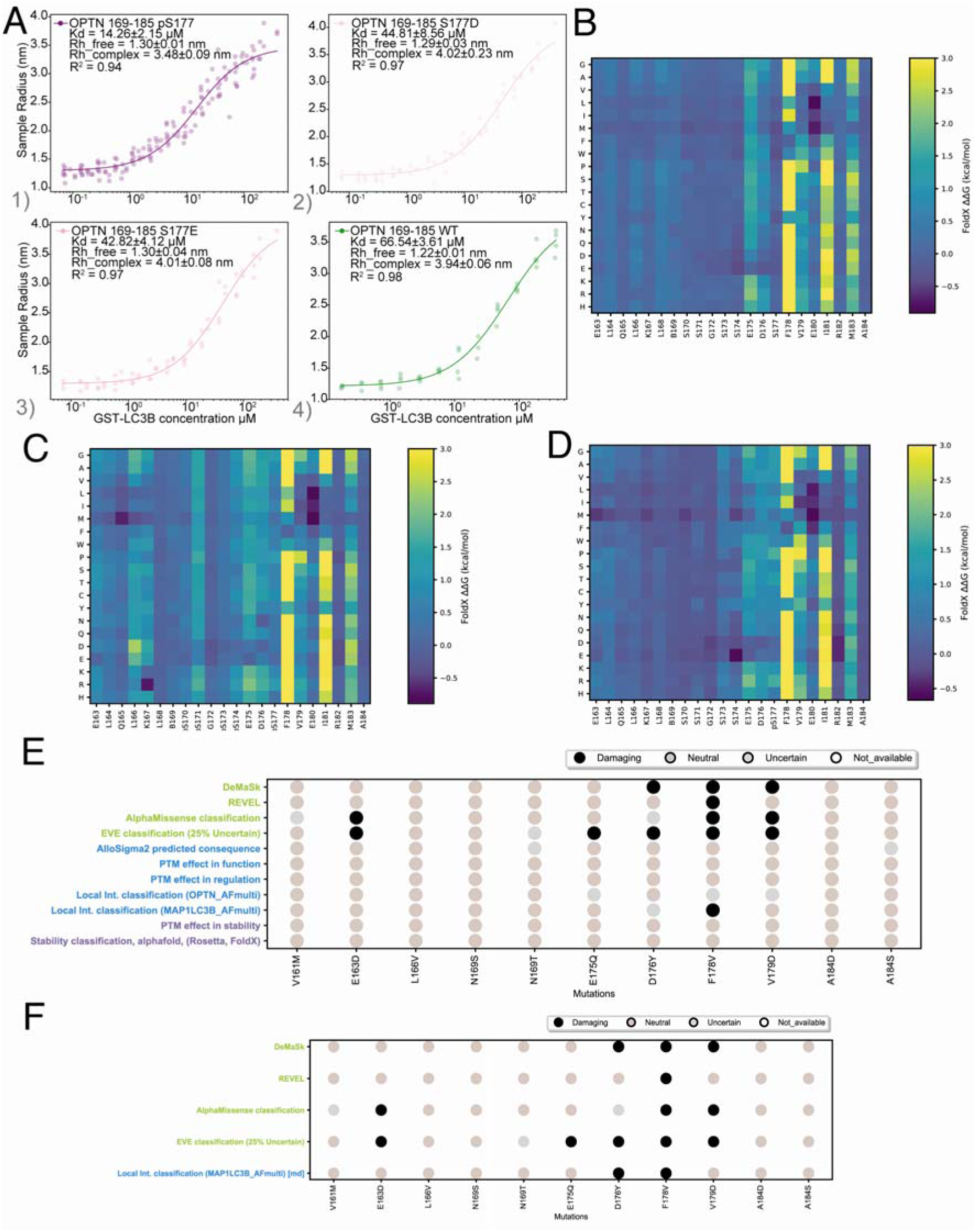
High-throughput mutational scans. (A) Binding isotherms of OPTN_169-185_ (1) pS177, (2) S177D, (3) S177E, and (4) WT FITC-labeled peptide variants to GST-LC3B by Microfluidic Diffusional Sizing. Raw data of all biological and technical replicates are plotted as dots, while the lines represent the non-linear fit curves. The fitted K_d_, Rh_free and Rh_complex values are reported as the global fit ± standard error across biological replicates. R^2^ values indicate the goodness of fit for each curve. (B-D) The heatmaps for the saturation scan in silico for LC3B_1-120_-OPTN_158-191_ (B), LC3B_1-120_-p5OPTN_158-191_ (C), and LC3B_1-120_-p1OPTN_158-191_ (D) are illustrated with a truncation of the ΔΔG values at 3 kcal/mol for visualization purposes. We can observe that the pentaphosphorylation activates binding hotspots in the N-terminal flanking region and decreases the importance of the negatively charged residues in the proximity of the core motif (E175 and D176) for the binding to LC3B (B-C). The effect cannot be recapitulated by the single phosphorylation (C-D). E-F) The dotplot summarized the results of the MAVISp assessment in the simple (E) and ensemble (F) modes. Black, white, and grey dots represent mutations predicted with a damaging, neutral, or uncertain effect for one of the MAVISp modules, respectively.

We had three main goals: i) evaluate if, with the mutational scan, we can predict the experimentally known stabilizing effect of the phosphorylations on the binding ^33,34^ to provide a baseline for similar future studies on other phosphorylatable LIRs; ii) to use the data to understand in more details the structural mechanisms promoted by the phosphorylation.

At first, we designed the calculations to evaluate the effect of introducing the phosphorylations on the structure of the unphosphorylated complex. In addition, we also used the structures from the phosphorylated complex and assessed the effect of removing the phosphorylation, which should destabilize the binding in the case of FoldX. We used this ‘reverse approach’ since FoldX has limitations in predicting stabilizing effects^97,98^. Thus, we speculate that it could be an effective alternative strategy to apply for the de novo discovery effects induced by phosphorylations on SLiMs. In more detail, the reverse approach entails a mutational scan with Foldx on ensembles of the phospho-variant, where the assessment is on the effect induced by substituting the mutation to the original unphosphorylated amino acid. In this kind of comparison, for example, if one observes only a marginal effect when a PTM/mutation is introduced starting from the wild-type ensemble of conformations but a clearly destabilizing effect when this PTM/mutation is removed in the ensemble of the modified variant, it can be more confidently predicted that the mutation is stabilizing the wild-type variant of the protein. In our comparison with experimental data, we also added the variant F178W of the OPTN LIR, which should mimic the interaction occurring for p62 with the HP1 pocket of LC3B.

At first, we observed that we could not recapitulate the stabilizing effects induced on the binding by the mutations F178W of the complex between LC3B and the unphosphorylated OPTN construct since the mutation was predicted with a neutral effect in most of the structures with regards to binding free energy. However, we observed, for some of the structures of the MD replicate2, a stabilizing effect induced by F178W with changes in binding ΔΔG of approximately -2 kcal/mol. F178W is of particular interest since it should mimic the interaction of the conventional p62 LIR using the wild-type ensemble of the complex.

When we assessed the effect of the phosphorylation starting from the unphosphorylated complex, we obtained average ΔΔGs values lower than 0, with pS177 and pS174 as the most important modifications to increase the binding affinity. Although the average differences were lower than the threshold of 1 kcal/mol generally used to pinpoint stabilizing or destabilizing variants for a protein-protein interaction. A group of conformations from the MD simulations featured values lower than the threshold, as observed for the F178W mutation introduced in the unphosphorylated variant. With the reverse approach, i.e., using the phospho-variant and measuring the effect of reintroducing a serine, we also observed a small average destabilizing effect (i.e., average ΔΔG values of 0.4-0.8 kcal/mol). The small changes might be due to the fact that the MutateX protocol only allows considering one mutation at a time^71^. This implies that the other phosphorylations were still included in the model when assessing a change from phospho-serine to serine at one site. We did not include calculations using multiple mutations of the phospho-sites simultaneously since empirical free energy functions, such as the one applied here with local sampling, are often not able to capture with accuracy the contribution of multiple mutations and would rather provide predictions that are the results of additive contributions from the single mutations.

Furthermore, we used the data from the saturation scan to identify residues that could contribute specifically to the binding affinity upon phosphorylation due to the rewiring of the interactions at the binding interface in the presence of phosphorylation (**Figure 4B-D**). To this goal, we selected residues that could serve as binding hotspots in the LIR (or LC3B) only in its pentaphosphorylated form and with marginal contributions in the unphosphorylated variant (**Table S5**, **Figure 5B-D**). We also verified if the mono-phosphorylation of S177 was sufficient to promote the same effects (**Table S5,** **Figure 5D**).

Apart from confirming the importance of the phosphorylation sites, we identified three sites in the N-terminal flaking region as key only for the phosphorylated variant, i.e., E163 (when mutated to S, G, T, H, or R), different mutations at the positions L166 (especially substitutions to I, M, F or R), and K167 when mutated to small, aromatic, or negatively charged residues. The single phosphorylation does not seem to be sufficient to exert the same N-terminal effects as the ones observed for the penta-phosphorylated variant (**Table S5**). On the other side, E175 seems to have a more central role in the interaction of the unphosphorylated variant than the penta-phosphorylated one, in agreement with the changes observed in salt-bridge interactions (**Figure 4B-D**). D176 and M183 are sensitive to changes in binding free energies upon mutation both in the unphosphorylated and phosphorylated variants.

We also noticed different amino acid preferences upon phosphorylation for the LC3B sites at the interface. For example, mutating D19 to aromatic or positively charged amino acids could further stabilize the complex with the pentaphosphorylated OPTN peptide. K49 and K51 are slightly stronger interaction hotspots for the phospho-peptide than the unphosphorylated variant. R37 is instead a centra hotspot only for the phosphorylated peptide. In fact, R37 can only tolerate substitutions to lysines in the LC3B-p5OPTN complex, in agreement with its importance in mediating the interaction strength and salt bridges with the N-terminal region of OPTN.

### Effects of disease-related mutations of OPTN on the binding with LC3B

The methodologies used above can also be applied to scrutinize the effects of disease-related variants, which still have uncertain significance or conflicting evidence and pose a challenge to biomedical research ^99^.

We identified twelve variants reported in cancer databases or in ClinVar (**Table 2**, **Figure 5E-F**) located in the OPTN LIR or its flanking regions (residues 158-191).

**Table 2.** Disease-related mutations and pathogenicity scores. If the source for the variant is ClinVar we reported the review status and the classification. DeMask generally classifies as loss-of-function variants with a fitness score < 0 and gain-of-function > 0. We used as a threshold for damaging variants a DeMask score < -0.3 or > 0.3 to decrease the number of false positives. A similar strategy is applied to the REVEL score.

| variant | source | REVEL | DeMask delta fitness | EVE | Alphamissense | $\Delta\Delta G$ binding - p5 | $\Delta\Delta G$ binding - p1 |
| --- | --- | --- | --- | --- | --- | --- | --- |
| V161M | ClinVar (1, VUS) | 0.435 | -0.201 | Benign | Ambiguous | Neutral | Neutral |
| E163D | ClinVar (1,VUS) | 0.404 | -0.264 | Pathogenic | Pathogenic | Neutral | Neutral |
| L166V | cBioPortal | 0.378 | -0.132 | Benign | Benign | Potentially Damaging (0.8-1.6 kcal/mol) | Neutral |
| N169S | cBioPortal | 0.161 | -0.010 | Benign | Benign | Neutral | Neutral |
| N169T | cBioPortal | 0.227 | -0.059 | Uncertain | Benign | Neutral | Neutral |
| E175Q | cBioPortal COSMIC | 0.309 | -0.092 | Pathogenic | Benign | Potentially Neutral (0.2-1.3 kcal/mol) | Damaging (1.0-145 kcal/mol) |
| D176Y | cBioPortal COSMIC | 0.442 | -0.317 | Pathogenic | Ambiguous | Potentially Neutral (0.2-1.2 kcal/mol) | Potentially Damaging (0.9-1.6 kcal/mol) |
| F178V | cBioPortal, COSMIC | 0.58 | -0.341 | Pathogenic | Pathogenic | Damaging (3.0-4.5 kcal/mol) | Damaging (3.0-3.5 kcal/mol) |
| V179D | ClinVar | 0.482 | -0.389 | Pathogenic | Pathogenic | Neutral | Neutral |
|  | (1,VUS) |  |  |  |  |  |  |
| A184D | cBioPortal | 0.245 | -0.091 | Benign | Benign | Neutral | Neutral |
| A184S | cBioPortal<br>COSMIC | 0.159 | 0.042 | Benign | Benign | Neutral | Neutral |

We applied a structure-based framework that allow to link predicted pathogenic effects with the underlying molecular mechanisms exerted by the mutation^100^.

None of the variants seem to affect OPTN homodimerization, and thus, potentially its oligomerization, even if E175Q, F178V, and V179D, remains uncertain according to the analyses (**Figure 5E**).

The F178V is predicted with pathogenic effects and the underlying mechanism is in relation to an abolishment of the interaction with LC3B (**Figure 5E-F**). The predicted pathogenic variant D176Y has a damaging effecto on the binding with LC3B for the unphosphorylated forms of OPTN using the ensemble mode of MAVISp, i.e., which accounts for protein dynamics (**Figure 5F** and **Table 2**). L166V seems to be damaging for the binding only for the phosphorylated form of the OPTN-LC3B complex, but its damaging effect cannot be recapitulated by the single phosphorylation at S177 (**Table 2**).

The different effects that disease-related mutations can exert depending on the phosphorylation status of the protein are interesting from a broader point of view. Approaches to predicting the effects of mutations often neglect the interplay between the mutation itself and the phosphorylation status of the target protein. Our results point out the need to design and implement a new module for the ensemble mode of the MAVISp framework, where the differential effects of mutations depending on a PTM can be captured. Of interest, V179D, V161M, and E163D are reported as a variant of uncertain significance in glaucoma and amyotrophic lateral sclerosis in ClinVar. Even though EVE and AlphaMissense predict a possible pathogenic effect for E163D and V179D (**Figure 5E-F**, **Table 2**), the structure-based assessment suggests that they have neutral effects. Instead, the MAVISp assessment and all the pathogenicity scores agree on the neutral effect of V161M, in agreement with what was observed in a population study for the same variant^101^.

### Microfluidic Diffusional Sizing can accurately estimate changes in Kd upon substitutions in the phosphorylated variant of OPTN

To further evaluate stabilizing or destabilizing effects of amino acid substitutions on the interaction between LC3B and phosphorylated optineurin, we established an approach based on Microfluidic Diffusional Sizing (MDS)^48,49,102^ to study the binding affinity between GST-tagged LC3B and different peptide variants corresponding to the 169-185 residues of the full-length OPTN. We have purchased FITC-labeled peptides and titrated them with increasing concentrations of GST-LC3B to follow changes in hydrodynamic radius that could be fitted to obtain a Kd value for each peptide. The low sample volume of GST-LC3B needed per binding curve (8 µL) allowed us to upconcentrate GST-LC3B to the hundreds of micromolar levels needed to saturate the binding signals for most peptide. Due to synthesis issues with penta-phosphorylated peptides, we introduced substitutions in the pS177 mono-phosphorylated peptide of OPTN. Luckily, we observed in the simulations and the free energy scans discussed above that many of the structural effects observed in the penta-phosphorylated variant are also recapitulated by the single phosphorylation. At first, we measured a Kd of 13.7 μM (**Figure 5A1**) for the mono-phosphorylated variant, in good agreement with what was previously shown with isothermal titration calorimetry^34^. Moreover, we added as a control the unphosphorylated peptide bearing the F178W substitution that mimics the p62 LIR and resulted in a Kd of 15.91 μM, in range with the pS177 variant (**Figure 6A**, p = 0.214) as previously reported^34^. An alanine substitution at position 178 (F178A) of OPTN abolishes the hydrophobic residue for the interaction with the HP1 pocket of LC3B. This variant combined with the phosphorylation at position 177 caused an increase of Kd of 5-fold (**Figure 6B**, p = 0.0019). This agrees with what we observed in the scans in silico where the introduction of F178A in the mono-phosphorylated construct resulted in average changes in binding ΔΔG of approximately 4.5 kcal/mol. This suggests that the phosphorylation alone cannot compensate for the loss of binding affinity caused by removing the central residue in the core motif. This result is also aligned with the possible damaging impact of the shortening of the side-chain length for the disease-related F178V variant discussed above (**Figure 5E-F**).

**Figure 6.**
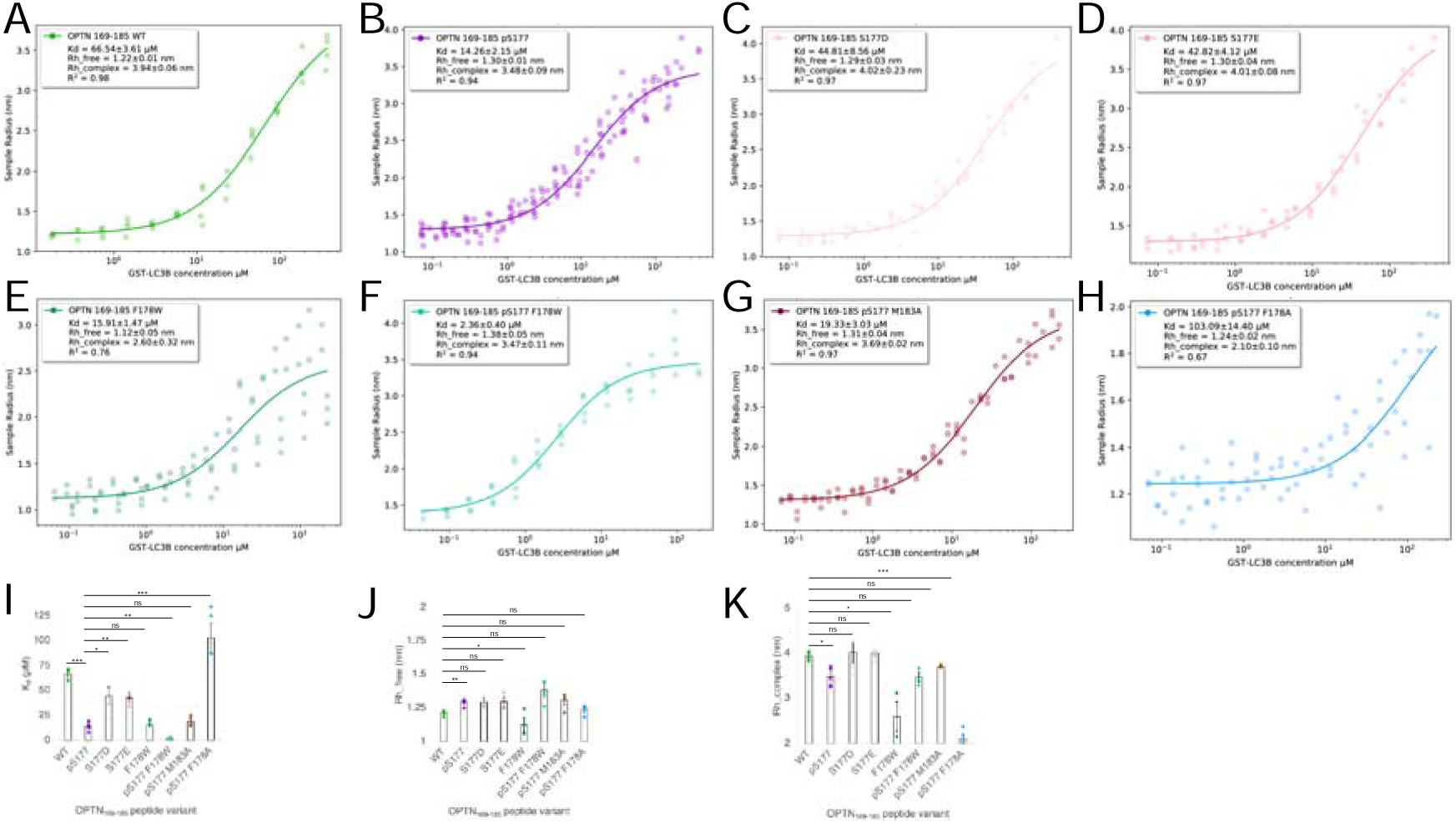
Equilibrium binding curves by Microfluidic Diffusional Sizing. (A-H) Binding isotherms of OPTN_169-185_ (A) WT, (B) pS177, (C) S177D, (D) S177D, (E) F178W, (F) pS177 F178W, (G) pS177 M183A, and (H) pS177 F178A FITC-labeled peptide variants to GST-LC3B by Microfluidic Diffusional Sizing. Raw data of all biological and technical replicates are plotted as dots, while the lines represent the non-linear fitting curves. The fitted K_d_, Rh_free and Rh_complex values are reported as the global fit ± standard error across biological replicates. R^2^ values indicate the goodness of fit for each curve. (I-K) Comparison of (I) K_d_, (J) Rh_free and (K) Rh_complex values across the tested peptide variants. Dots represent the fitted value for each biological replicate, while columns represent the global fit ± standard error across biological replicates. P values are generated through t-tests or t-tests with Welch’s correction, as described in Methods, to assess differences to the or WT or pS177 variant, and they are symbolized with “ns” if P > 0.05, “*” if P ≤ 0.05, “**” P ≤ 0.01, and “***” if P ≤ 0.001.

On the contrary, the F178W substitution that mimics a W-type LIR combined with the pS177 phosphorylation can increase the binding affinity 7-fold compared to the phosphorylation alone (**Figure 6C**, p = 0.003). To evaluate the effects of variants in the LIR flanking region, as well as to test the limit of the MDS approach in capturing smaller changes in binding affinity, we assessed the M183A substitution, which is predicted to destabilize the interaction for both the phosphorylated and unphosphorylated variants of OPTN with approximately 2 kcal/mol. We observed similar Kd values within the experimental error, from combining the phosphorylation at S177 and M183A compared to the phosphorylation alone (**Figure 6D, p = 0.25**). Our results collectively suggest that the MDS binding assay could be used to identify and validate predicted changes in binding affinity of more than 2 kcal/mol. Therefore, we did not work experimentally with the variants reported in **Table 2 or S4**, since all the predicted average changes for mutations in residues outside the core motif were mostly 1-2.5 kcal/mol (**Table S5**).

### Kinetics of OPTN LIR motif interaction by Surface Plasmon Resonance

We then provided quantitative kinetic constants for the interaction between the OPTN LIR and its partner LC3B. Traditionally, measuring these interactions has proven to be challenging due to the rapid on-off kinetics characteristic of LIR motifs—so rapid, in fact, that literature reports are scarce to non-existent^103,104^.

Our strategy involved immobilizing GST-LC3B onto a CM5 sensor chip and introducing increasing concentrations of LIR peptides. The sensorgrams generated were then modeled to a 1:1 binding interaction, from which we derived the association (k_on_) and dissociation (k_off_) rates (**Figure 7A-F**). Given that the k_off_ values exceeded the measurement capabilities of the Biacore X100 instrument, the fits are acknowledged as suboptimal. Conversely, most k_on_ values are on the brink of the instrument’s detection limit, yet they appear to be distinctly determinable. Therefore, we leaned on the steady-state K_d_ values, corroborated by MDS measurements, to extrapolate k_off_ rates. Despite the apparent stability issues with GST-LC3B on the chip that precluded multiple measurements, our analysis yielded meaningful insights. The sensorgram analysis of the OPTN_169-185_ unphosphorylated peptide revealed a steady-state K_d_ of 63.5 µM, consistent with MDS-derived and previously published values.^33,34^ The k_on_ was measured at 0.55 x 10^4^ M^-1^s^-1^, and with the steady-state K_d_, the k_off_ was computed at 3.51 x 10^-2^ s^-1^. Interestingly, while the off-rate aligns with the affinity of the high-affinity K1 peptide for GABARAP identified through phage-display^105^, the µM affinity of the natural OPTN LIR peptide is primarily modulated by a significantly reduced on-rate.

**Figure 7.**
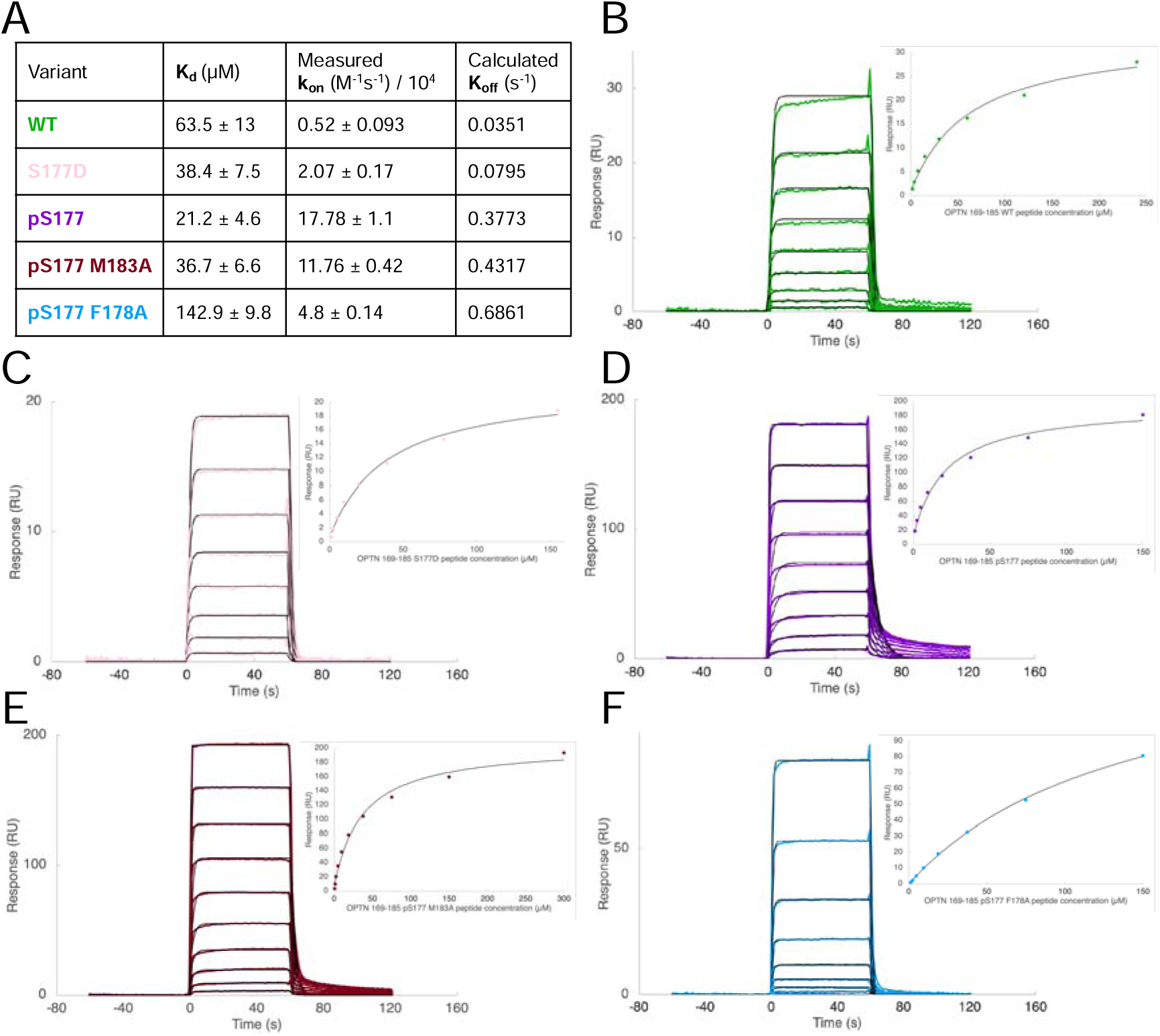
Kinetics of binding between the LIR motif of optineurin and GST-LC3B by Surface Plasmon Resonance. (A) Summary of the steady-state dissociation constants (K_d_) and on-rates (k_on_) for the five peptide variants tested: OPTN_169-185_ WT, S177D, pS177, pS177 M183A, and pS177 F178A. The off-rates (k_off_) were calculated from these values. The averages and standard errors were derived from non-linear regression fits of the binding curves. (B-F) Representative SPR sensorgrams showing the binding interactions of increasing concentrations of OPTN169-185 peptide variants (B) WT, (C) S177D, (D) pS177, (E) pS177 M183A, and (F) pS177 F178A with amine-coupled GST-LC3B on a CM5 chip. The colored traces represent experimental data, while the black lines indicate the kinetic fits based on a 1:1 binding model. Insets display the steady-state resonance values taken 10 seconds before the end of injection, collected in 5-second intervals, and plotted against peptide concentration to calculate steady-state Kd values (black line fits).

Introducing a phosphorylation at position 177 (pS177) in the OPTN_169-185_ peptide notably enhances the interaction dynamics with LC3B. This modification leads to a ∼30-fold increase in the on-rate to 17.7 x 10^4^ M^-1^s^-1^ and about a 10-fold increase in the off-rate to 37.7 x 10^-2^ s^-1^. However, the net effect is a 3-fold improvement in steady-state affinity to 21.2 µM due to the more pronounced increase in the on-rate. This dynamic shift is also observed, albeit to a lesser degree, with the phosphomimetic S177D substitution, leading to a steady-state affinity of 36.7 µM—improved from the wild type but not as potent as the true phosphorylation. When evaluating the pS177 peptide, substitutions in the flanking region (M183A) and core LIR motif (F178A) had detrimental effects on the steady-state affinity, impacting both kinetic rates. The M183A substitution led to nearly a 2-fold drop in affinity to 36.7 µM. However, limitations in data reproducibility prevent a robust significance assessment. The F178A substitution within the pS177 peptide, on the other hand, induced a marked decrease in affinity, with a K_d_ exceeding 100 µM, echoing the MDS findings and underscoring that phosphorylation cannot compensate for the loss of the critical HP1 pocket aromatic residue. In conclusion, our study not only fills a gap in the current understanding of LIR motif dynamics but also establishes a reliable experimental framework for future investigations into the kinetics of these transient but pivotal interactions.

## Conclusions

Phosphorylation can modulate LIR motifs and compensate for the lower binding affinity in LIRs. Phosphorylatable sites have been reported and experimentally verified for other LIR motifs with an impact on binding affinity and recognition of LC3/GABARAP proteins^19^. It is thus of fundamental importance to investigate the role of these phosphorylations on the structure and function of LIR-containing proteins and their complexes.

We here employed the OPTN LIR as a model system of phospho-switches for the LIR-ATG8 recognition, thanks to the availability of biophysical data on its phosphorylations and mutations that can be used to compare and validate our simulation data^33,34^. Moreover, we generated new experimental data on different OPTN variants applying for the first time a technique based on Microfluidic Diffusional Sizing to protein-SLiM interactions and measurements based on Surface Plasmon Resonance. In addition, we shed light on the role of the flanking-region N-terminal to the core LIR motif of OPTN as a sensor of long-range effects induced by OPTN phosphorylation.

Our result, together with previous results on another LIR-containing protein^94^, show that phosphorylation can exert this effect by impacting the flexibility of the entire LIR peptide once the complex is formed. Moreover, the post-translational modification can induce long-range effects stabilizing the electrostatic interactions in the flanking region of the peptide, rewiring the network of intermolecular interactions. This rewiring also causes changes in hotspots for binding in the flanking regions and their contribution to the binding with LC3B. The phosphorylation of the residue pS177 alone seems to be sufficient to promote changes in solvent accessibility for the LC3B residues involved in interactions with the central part of the OPTN LIR motif. This is mirrored by the fact that the phosphorylations of the LIR cannot rescue the flipping movement of the OPTN phenylalanine residue in the HP1 binding pocket. Furthermore, the multiple phosphorylation can further stabilize the complex expanding the β-sheet structure in the binding cleft. On the other hand, the mono-phosphorylation is not sufficient to decrease the protein flexibility of OPTN in the complex and cannot elicit long-range effects to recruit residues from the N-terminal flanking regions into the complex formation.

Phosphorylation of OPTN is important to regulate its cellular function and the structural changes induced by it should also be considered in the context of understanding the impact of disease-related mutations. The approach used in this study to investigate the effects of disease-related variants on the phosphorylated form of the protein can serve to design future studies on other disease-related targets for which phosphorylation events have been experimentally characterized.

Overall, our comprehensive investigation into the phospho-modulation and flanking region contributions of LIR motifs, using OPTN as a model system, provides valuable insights into the intricate interplay between phosphorylation, structural alterations, and functional implications in LIR-containing proteins. The elucidation of long-range effects, alterations in binding hotspots, and the impact on solvent accessibility not only enhances our understanding of the regulatory role of phosphorylation in OPTN but also underscores its significance in the broader context of disease-related mutations. Our methodology, encompassing biophysical data, simulations, and innovative experimental techniques, establishes a robust framework for future studies on diverse targets, contributing to a deeper comprehension of the cellular processes governed by phospho-regulated Short Linear Motifs.

## Author Contributions (CRediT Classification)

Conceputalization: EP. Data Curation: EP, MU, OA, VS, ML. Formal Analysis: MU, OA, EP Funding Acquisition: EP. Investigation: EP, OA, MU, VS, ML. Methodology: OA, EM, ML, MT. Project administration: EP. Resources: EP. Software: MT, ML, VS, MU, EP. Supervision: EP, ML, EM. Validation: MU, ML, EP. Visualization: MU, ML, OA, VS, EP. Writing – Original Draft: EP, OA, MU, VS, ML, EM. Writing – Review and Editing: All the coauthors.

## Acknowledgments

Our research has been supported by Danmarks Grundforskningsfond (DNRF125), Carlsberg Foundation Distinguished Fellowship (CF18-0314), Hartmanns Fond (R241-A33877) and NovoNordisk Fonden Bioscience and Basic Biomedicine (NNF20OC0065262) to EP group. Part of the calculations have been supported by a ISCRA CINECA Grant, DECI 16h and a EuroHPC Regular Access Grant (EHPC-REG-2023R01-051) on Discoverer. The calculations described in this paper were performed using the DeiC National Life Science Supercomputer at DTU. The funders had no role in study design, data collection or analysis, decision to publish, or manuscript preparation.

We would like to thank Kirsten Frederisken for statistical advice, as well as Burcu Aykac Fas, Mukesh Kumar, and Željka Sanader Maršić for their involvement in the early stages of this project.

