## Supplementary figures and images for "Decoding Phospho-Regulation and Flanking Regions in Autophagy-Associated Short Linear Motifs: A Case Study of Optineurin-LC3B Interaction"

### Figure S1

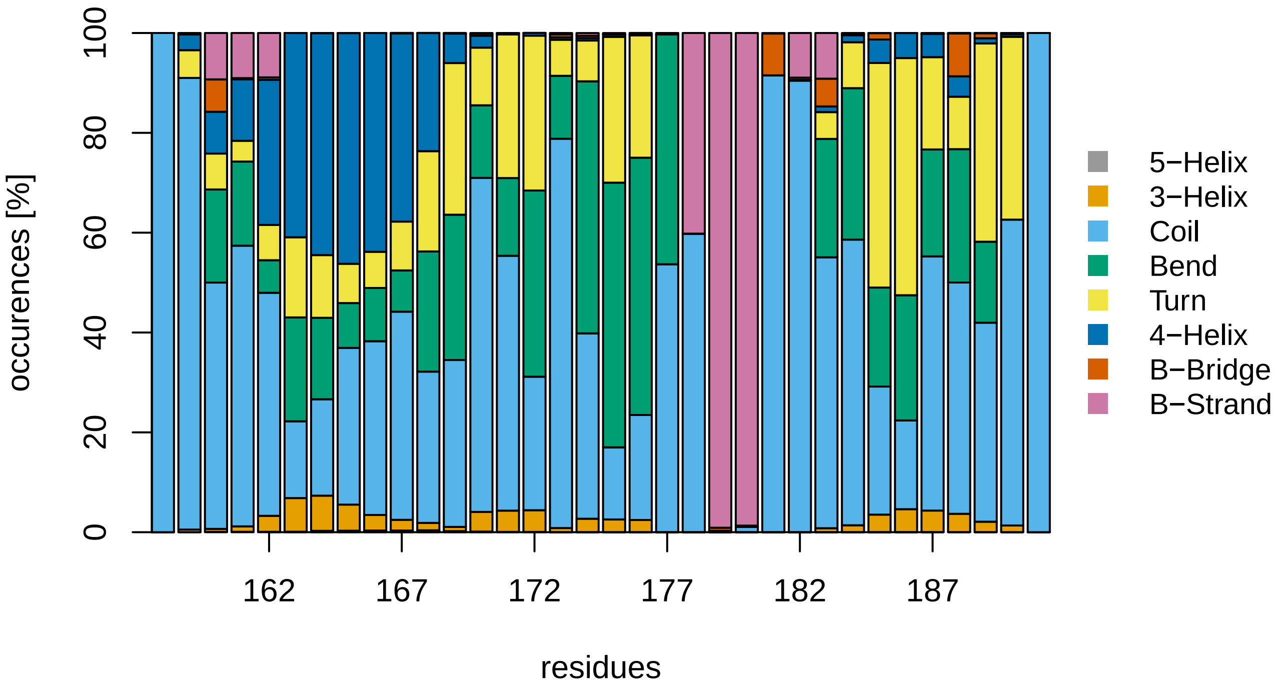
