## Supplementary material for "Decoding Phospho-Regulation and Flanking Regions in Autophagy-Associated Short Linear Motifs: A Case Study of Optineurin-LC3B Interaction": Figure S2

**Figure S2. Solventaccessible surface for F178 (i.e., the residue to bind the HP1 pocket of LC3B) of OPTN in the MD simulations.**

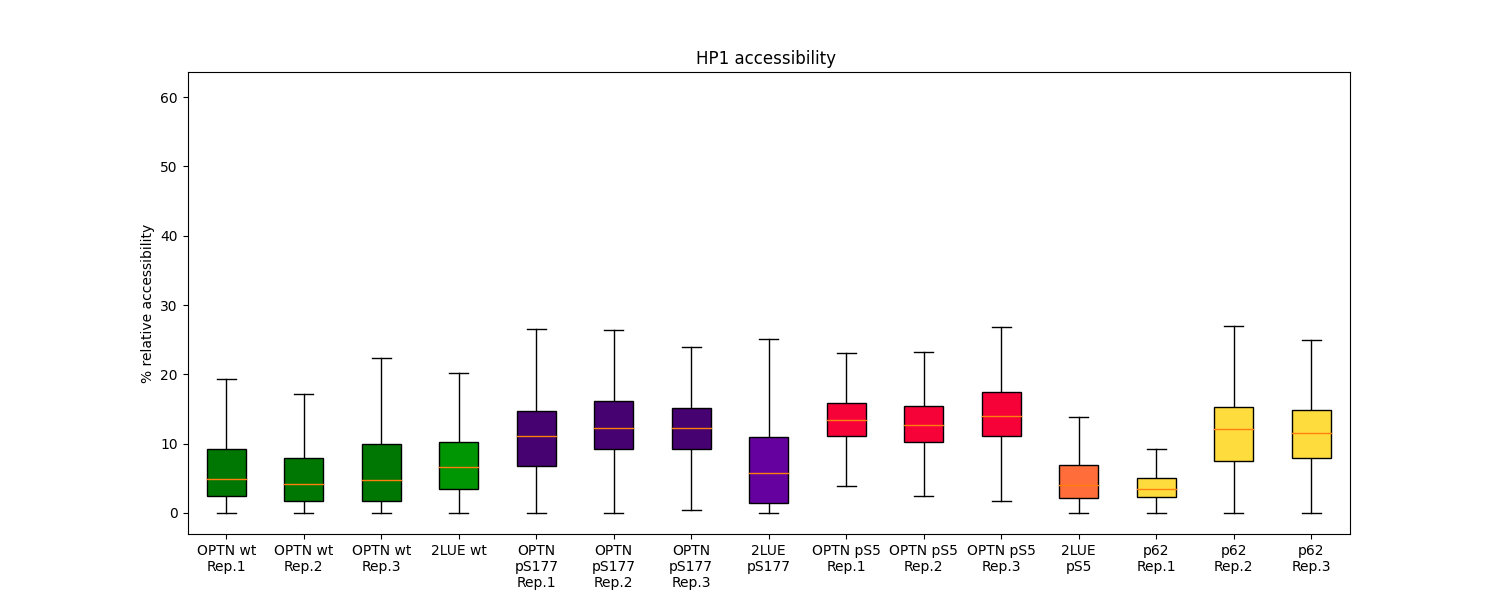
