## Supplementary material for "Decoding Phospho-Regulation and Flanking Regions in Autophagy-Associated Short Linear Motifs: A Case Study of Optineurin-LC3B Interaction": Figure S3

**Figure S3. Solvent-accessible surface for I181 (i.e., the residue to bind the HP2 pocket of LC3B) of OPTN in the MD simulations.**

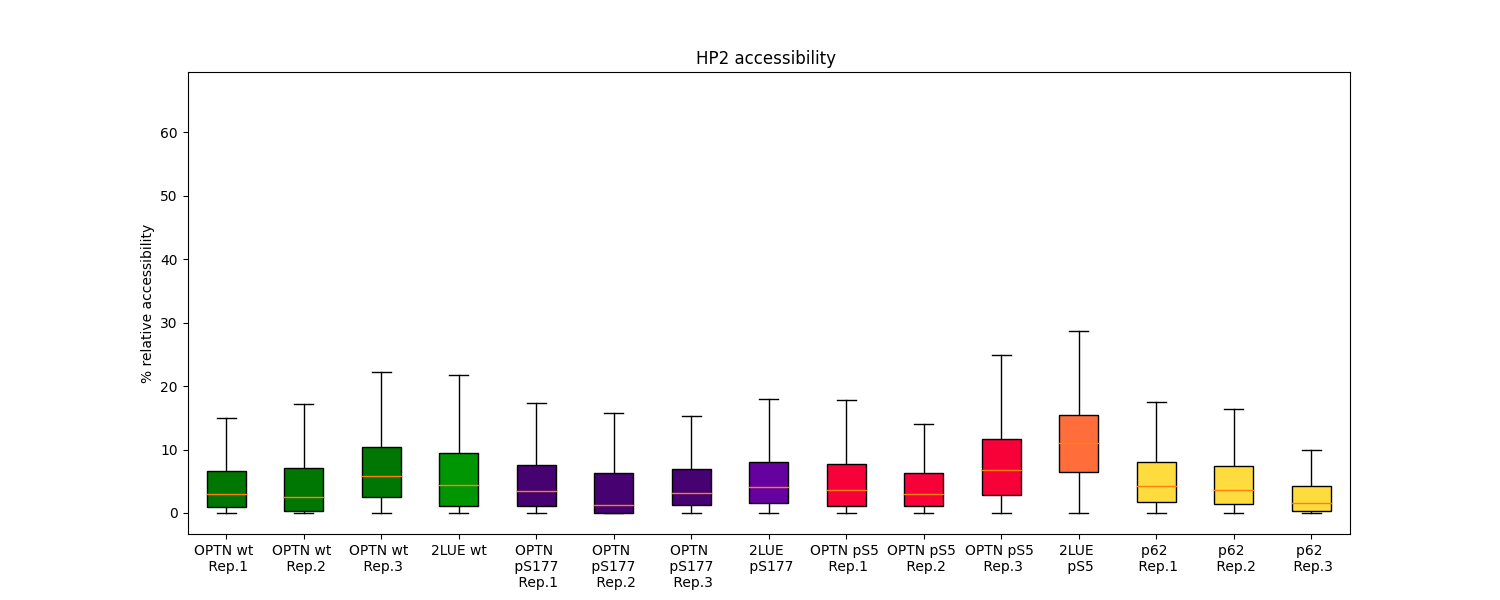
