## Supplementary material for "Decoding Phospho-Regulation and Flanking Regions in Autophagy-Associated Short Linear Motifs: A Case Study of Optineurin-LC3B Interaction": Table S1

**Table S1. Summary of simulations used in the study**

| **System** | **Force Field** | **Capping LC3B** | **Capping LIR** | **Sampling Method** | **Length (Time)** | **N° replicates** |
| --- | --- | --- | --- | --- | --- | --- |
| LC3B_1-119_-p5OPTN_168-185_ | CHARMM22* | NH2-COOH | NH2-COOH | unbiased MD | 1 µs | 4 |
| LC3B_1-120_-OPTN_158-191_ | CHARMM22* | NH3^+^-COOH | NH2-COOH | unbiased MD | 1 µs | 3 |
| LC3B_1-120_-OPTN_158-191_ | CHARMM22* | NH3^+^-COOH | NH2-COOH | parallel bias metadynamics | 0.5/0.6 µs | 2 (two different sets of CVs) |
| LC3B_1-120_-p1OPTN_158-191_ | CHARMM22* | NH3^+^-COOH | NH2-COOH | unbiased MD | 1 µs | 3 |
| LC3B_1-120_-p1OPTN_158-191_ | CHARMM22* | NH3^+^-COOH | NH2-COOH | parallel bias metadynamics | 0.5 µs | 1 |
| LC3B_1-120_-p5OPTN_158-191_ | CHARMM22* | NH3^+^-COOH | NH2-COOH | unbiased MD | 1 µs | 3 |
| LC3B_1-120_-p5OPTN_158-191_ | CHARMM22* | NH3^+^-COOH | NH2-COOH | parallel bias metadynamics | 0.5/0.6 µs | 1 |
| LC3B_1-120_-OPTN_169-185_ | CHARMM22* | NH3^+^-COOH | NH2-COOH | unbiased MD | 1 µs | 1 |
| LC3B_1-120_-p1OPTN_169-185_ | CHARMM22* | NH3^+^-COOH | NH2-COOH | unbiased MD | 1 µs | 1 |
| LC3B_1-120_-p5OPTN_169-185_ | CHARMM22* | NH3^+^-COOH | NH2-COOH | unbiased MD | 1 µs | 1 |
